# Project ChemicalBlooms: Collaborating with Citizen Scientists to Survey the Chemical Diversity and Phylogenetic Distribution of Plant Epicuticular Wax Blooms

**DOI:** 10.1101/2023.11.11.566687

**Authors:** Dien Nguyen, Nicole Groth, Kylie Mondloch, Edgar B. Cahoon, Keith Jones, Lucas Busta

## Abstract

Plants use chemistry to overcome diverse challenges. A particularly striking chemical trait that some plants possess is the ability to synthesize massive amounts of epicuticular wax that accumulates on the plant’s surfaces as a white coating visible to the naked eye. The ability to synthesize basic wax molecules appears to be shared among virtually all land plants and our knowledge of ubiquitous wax compound synthesis is reasonably advanced. However, the ability to synthesize thick layers of visible epicuticular crystals (“wax blooms”) is restricted to specific lineages and our knowledge of how wax blooms differ from ubiquitous wax layers is less developed. Here, we recruited the help of citizen scientists and middle school students to survey the wax bloom chemistry of 78 species spanning dicot, monocot, and gymnosperm lineages. Using gas chromatography-mass spectrometry, we found that the major wax classes reported from bulk wax mixtures can be present in wax bloom crystals, with fatty acids, fatty alcohols, and alkanes being present in many species’ bloom crystals. In contrast, other compounds including aldehydes, ketones, secondary alcohols, and triterpenoids were present in only a few species’ wax bloom crystals. By mapping the 78 wax bloom chemical profiles onto a phylogeny and using phylogenetic comparative analyses, we found that secondary alcohol and triterpenoid-rich wax blooms were present in lineage-specific patterns that would not be expected to arise by chance. That finding is consistent with reports that secondary alcohol biosynthesis enzymes are found only in certain lineages, but was a surprise for triterpenoids, which are intracellular components in virtually all plant lineages. Thus, our data suggest that a lineage-specific mechanism other than biosynthesis exists that enables select species to generate triterpenoid-rich surface wax crystals. Overall, our study outlines a general mode in which research scientists can collaborate with citizen scientists as well as middle and high school classrooms not only to enhance data collection and generate testable hypotheses, but also directly involve classrooms in the scientific process and inspire future STEM workers.

**Significance Statement:** Plants coat themselves in a protective layers of waxes. Some plants produce exceptionally thick layers of wax (“wax blooms”), and these thick layers are associated with enhanced abilities to tolerate stress, including drought and insect attack. In collaboration with citizen scientists and middle school classrooms, we provide an overview of the chemistry and phylogenetic distribution of plant epicuticular wax blooms. These data constitute a foundation upon which future studies of diverse wax blooms and their functions can build.

## 1. Introduction

Plants use a diverse array of chemical compounds to solve challenges they face. Some plant chemicals are produced by virtually all plant species (core metabolites) while other compounds are produced only by select species (lineage-specific metabolites), thus defining the extremes of a spectrum onto which plant chemical products can be placed. To deal with the challenge of living out of water, virtually all land plants coat themselves in a mixture of wax chemicals that prevent non-stomatal water loss. This water barrier consists of two layers of waxes: the intracuticular wax layer, which is embedded into a polyester scaffold called cutin, and the overlaying epicuticular wax layer, which consists of waxes in amorphous or crystalline form. The specific wax chemicals that comprise these two layers can be mixtures of core metabolites (for example, fatty acids, fatty alcohols, fatty aldehydes, alkanes, and alkyl esters) or lineage-specific metabolites (for example, secondary alcohols, ketones, and triterpenoids). Studies have shown that the composition of the two wax layers can be distinct from one another (1), and can vary between plant species, tissues of the same species, and even cell types of a given plant tissue (2–4). Mounting evidence suggests that the intracuticular layer creates the water barrier (5–8), while the epicuticular layer performs other functions, including protecting against UV radiation (9), mediating plant-insect interactions (10–13), plant-microbe interactions (14, 15), and a plant’s self-cleaning ability (16, 17). All of these functions, including water retention, are influenced by the chemical composition of the two wax mixtures (5, 18–21). Thus, by improving our understanding of plant wax chemistry, we can help advance research and crop improvement efforts related to a diverse array of plant functions–a critical pursuit in the face of a changing climate.

In some plant species’ epidermal cells, wax biosynthetic processes are so active that epicuticular waxes accumulate visibly as a white coating, often referred to as a “wax bloom”.

These coatings are so thick that they can be seen with the naked eye and removed with a brush or finger (Fig. 1). Wax blooms have been studied on a variety of crops. For example, a field study with *Triticum aestivum* (wheat) found the presence of wax blooms to be positively correlated with both cooler canopy temperature and grain yield under water-deficient conditions (22). A sorghum field study found that, under water-deficient conditions, water became a limiting factor to plants with lower wax bloom load sooner than it did to plants with a higher wax load (23). In other crops, such as some *Vaccinium* species (blueberry), wax blooms play a role in preventing fruit desiccation (24). Wax blooms can also be a desirable consumer trait and play a role in post-harvest storage (25, 26). Thus, experimental evidence from diverse crops shows that wax blooms play roles in stress tolerance, post-harvest biology, and consumer perception. These roles emphasize the importance of advancing our general understanding of epicuticular wax blooms.

**Fig. 1.**
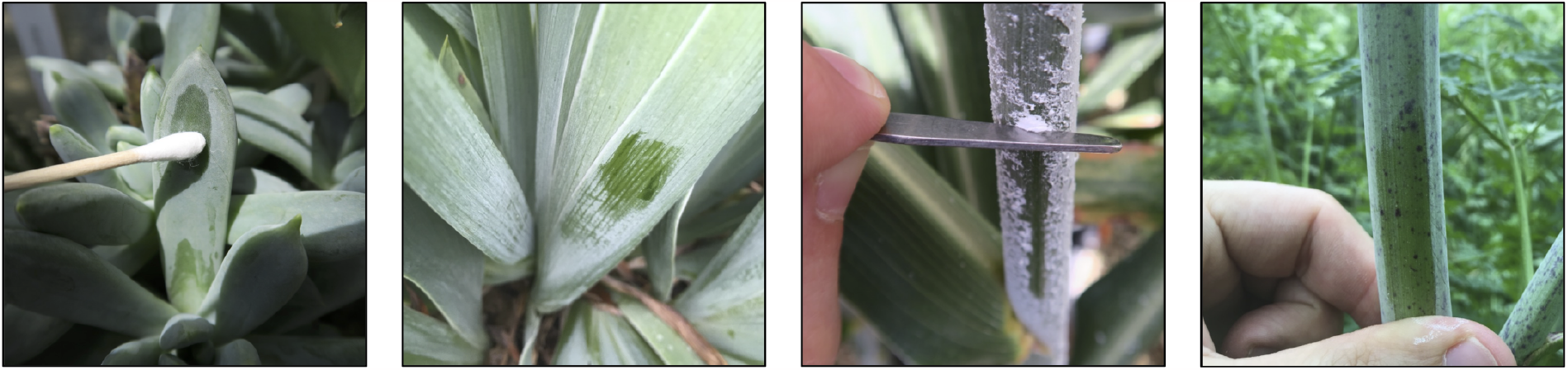
Photographs of epicuticular wax blooms. Representative photographs of epicuticular wax blooms on, from left to right, *Graptopetalum paraguayense, Iris germanica, Sorghum bicolor*, and *Daucus carota*. In each photograph, the wax bloom has been partially removed to highlight the thickness of the epicuticular wax layer and its ease of removal.

Two open questions regarding plant epicuticular wax blooms are (i) whether and how the chemistries of the wax bloom crystals are significantly different from the chemistries of plant wax mixtures that have been described so far, and (ii) which plant species are capable of making epicuticular wax blooms and how are those species distributed across the plant phylogeny. To facilitate future systematic investigations of plant epicuticular wax blooms, this project aimed to provide data and analyses to answer these two questions. Not all plant species produce epicuticular wax blooms. Therefore, searching out a diversity of bloom-producing species for study would normally require considerable time and long travel distances. For these reasons, we used social media to recruit the aid of citizen scientists. We sent participating citizen scientists bio-prospecting kits with which to collect samples of wax chemical blooms that could be returned to our laboratory by mail for subsequent chemical analysis. Here, with the help of more than 70 citizen scientists, we provide a phylogenetically-resolved map of epicuticular wax bloom chemistry. We also apply phylogenetic comparative analyses to investigate the nature of lineage-specific compounds that occur in plant epicuticular wax blooms.

## 2. Results and Discussion

The objective of this study was to characterize the chemical composition and phylogenetic distribution of plant epicuticular wax blooms. To sample a broad number of plant species, we first worked with citizen scientists to collect samples for gas chromatography-mass spectrometry analysis (section 2A). We then mapped the chemical profiles onto a phylogenetic tree and used phylogenetic comparative analyses to examine patterns in the distribution of the chemical profiles across the phylogeny (section 2B).

### A. Chemistry of plant epicuticular wax blooms

To collect epicuticular wax blooms isolates from a wide range of plant species, we used social media to connect with citizen scientists, then sent the citizen scientists bioprospecting kits with which to collect isolates. The kits included (i) documentation on the significance of the project, (ii) instructions on how to collect plant identification information and wax bloom chemical isolates using the enclosed sample information sheet and cotton swabs, as well as (iii) a return envelope with which to mail the isolates back to our laboratory (Fig. 2A). Over the course of three years, we communicated with and were sent isolates and photographs by more than 70 citizen scientists. Collectively, they sent a total of 228 samples. We found that working with middle school and high school classrooms and scout troops was particularly effective because the teacher or group leader could help ensure that a large number of samples were collected, that plant identification information was documented as well as possible, and that isolates and information returned safely to our laboratory.

**Fig. 2.**
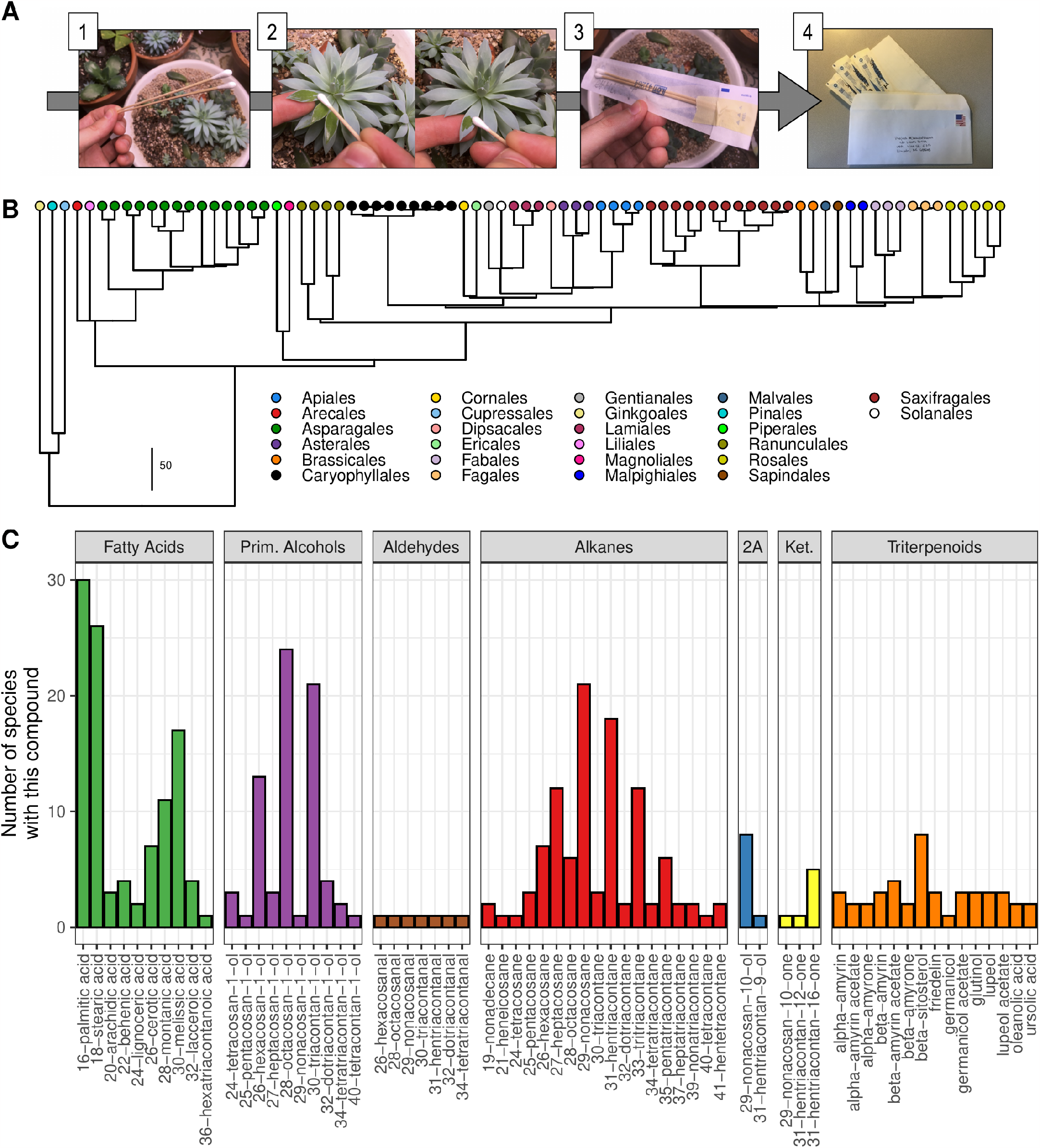
Prevalence of specific wax chemicals in epicuticular wax blooms. A. Schematic of the procedure used by citizen scientists to collect epicuticular wax bloom samples and send them to the analysis laboratory. **B**. Phylogeny of the 78 species analyzed in this study. This phylogeny was derived via pruning from a previously published megaphylogeney (27). Tips are color-coded according to plant families as indicated in the legend. The scale indicates time corresponding to branch length in millions of years. **C**. Bar chart showing the frequency with which each wax compound was found as a component of the epicuticular wax chemical bloom. The total number of blooms sampled was 78, so, for example, 16-palmitic acid was found in slighly fewer than half of the wax blooms sampled. Abbreviations: 2A = Secondary alcohols, Ket. = Ketones. The chain length of each aliphatic compound is given in front of the compound’s name on the x-axis.

Middle school students in Mr. Keith Jones’ class at White Rock Elementary in McDonald County, Missouri were recurring participants in our chemical bloom sample collection efforts. Mr. Jones noted that by involving students in this citizen science project, student engagement in related curricular activities was very high. Even a year later, students remembered details about waxes and their roles in plant defense and mediating the plant’s interactions with the environment. Mr. Jones reported that the students readily recognized that their work was a legitimate part of scientific research and that their work has gone toward increasing humanity’s understanding of the world around them. He also noted that many students, particularly young women, were clearly more excited about STEM after participating in the project. We acknowledge that these observations are anecdotal, but they are consistent with well-documented effects of active learning approaches to teaching. This consistency suggests that other bioprospecting-like plant chemistry projects will likely be good tools for inspiring the next generation of STEM professionals.

Next, we prepared a gas chromatography-mass spectrometry (GC-MS) sample from each wax bloom isolate we received from citizen scientists. Wax compound concentrations in the samples varied considerably, so appropriate dilution levels and injection volumes needed to be determined on a per-sample basis. In some cases, one or more circumstances prevented us from including a particular isolate in our final data set. Reasons for exclusion included (i) the amount of wax collected was apparently not sufficient for GC-MS analysis or (ii) the provided plant identification information was not sufficient to identify the plant at the genus level. Additionally, there were rare instances where multiple isolates from the same taxon displayed vastly different chemical profiles. Some of these variations were clearly due to contaminants, like phthalates. However, when the profiles appeared uncontaminated yet were significantly different, it indicated other potential variations like the plant’s age when sample, the season of collection, or the tissue from which the sample was taken. These factors can affect wax chemical compositions. Consequently, such taxa were left out of our final data set, highlighting a need for more in-depth analysis of these species before their wax chemistry should be reported. Even so, we eventually obtained quality GC-MS data from a total of 78 species spanning 26 plant families, with representatives from the dicot, monocot, and gymnosperm lineages (Fig. 2B). Thus, while approximately two-thirds of the isolates we received did not contribute to our final data set, the efforts of participating citizen scientists enabled the collection of quality data from species that spanned a considerable portion of the plant phylogeny.

Using the data from the 78 species described above, we first examined the types of wax compounds present in epicuticular wax blooms. Across the 78 species, we identified a total of 70 different wax compounds that belonged to seven major wax compound classes. Specifically, we found 10 fatty acids, 10 fatty alcohols, 7 fatty aldehydes, 18 alkanes, 7 ketones, 2 secondary alcohols, and 16 triterpenoids (Fig. 2C). Thus, it seems that the major wax classes that have been reported previously from bulk plant wax mixtures (epicuticular and intracuticular layers combined) can be present in the crystals of epicuticular wax blooms. We note that our experiments cannot rule out the existence of wax blooms comprised of alkyl esters due to the difficulties associated with detecting these high molecular weight compounds with split/splitless GC injectors. The seven epicuticular wax bloom compound classes were distinguished from one another by their diversity, with some classes containing a large number of members (alkanes and triterpenoids) and some classes only having a few members (aldehydes, ketones, and secondary alcohols). Among the compounds in the seven classes, some were much more common (i.e. found in a higher number of species) than others. C16 and C18 fatty acids, as well as C30 fatty acids, were found in the epicuticular wax blooms of >25 and >15 plant species, respectively (Fig. 2C). Alcohols with even total carbon numbers C26-C30 and alkanes with odd total carbon numbers C27-C33 were also found on many species (>10). In contrast, each of the aldehyde, ketone, secondary alcohol, and triterpenoid compound was found on fewer than ten species and many of these compounds were found on just one or two species. Based on our finding that some types of wax chemical blooms are very rare while others are quite common, we became interested in exploring the phylogenetic distribution of epicuticular wax bloom chemistry.

### B. Phylogenetic distribution of plant epicuticular wax bloom chemical profiles

Our next objective was to map the plant epicuticular wax bloom chemical profiles from section 2A onto a plant phylogeny. For this, we pruned a previously published plant megaphylogeny (27) so that it contained only tips representing species for which we had chemical data. We then plotted the resulting tree alongside a heat map depicting the abundance of each epicuticular wax chemical (Supplemental Fig. S1). We also plotted the tree alongside a horizontal bar chart showing the relative abundance of each compound class in each species (Fig. 3). The compound class-level plot revealed several interesting patterns. First, we noticed that some compound classes, including fatty acids, alcohols, and alkanes, were found in a high number of disparate lineages. This pattern of distribution is consistent with previous findings that suggest the pathways to these compound classes are ancient and are likely found in virtually all land plant lineages. In contrast to the widely-distributed compound classes, we observed that some compound classes were only found in a small number of species or were restricted to specific lineages. These lineage-specific wax bloom compounds included aldehydes, secondary alcohols, ketones, and triterpenoids. Some of these classes occurred in lineage specific clusters, for example, secondary alcohols in the gymnosperm genera *Cupressus, Picea*, and *Ginkgo*; secondary alcohols in the Ranunculales genera *Sanguinaria* and *Papaver*; triterpenoids in the Lamiales genera *Salvia, Monarda*, and *Lavandula*; and triterpenoids in the Saxifragales genera *Kalanchoe, Bryophyllum, Echeveria*, and *Hylotelephium* (Fig. 3A). This clustering suggested that a negative correlation might exist between chemical similarity and the phylogenetic distance separating species. In other words, the wax bloom chemistries of two closely related species are significantly more likely to be more similar than the chemistries of two distantly related species. To test this hypothesis, we computed and regressed the presence-absence similarity and quantitative similarity of the chemical profiles for each pair of species against the phylogenetic distance between each pair of species (Fig. 3B and C). We found a statistically significant negative correlation in both cases (p-value for slope < 0.05), suggesting that ecological or evolutionary factors are influencing plant epicuticular wax bloom chemical profiles, and that these compound classes were not distributed randomly over the phylogenetic tree. Based on that finding, we further hypothesized that the apparently clustered classes (aldehdyes, secondary alcohols, ketones, and triterpenoids) were found more frequently in closely related species than would be expected by random chance. To test that hypothesis, we computed the phylogenetic signal for each compound class and found that secondary alcohols and triterpenoids were indeed more tightly clustered together than would be expected by chance (lambda > 0.99, bonferroni corrected p-value for lambda < 0.001). Our analyses did not suggest that any other compound classes were distributed in a way that would not be expected by random chance. Nevertheless, ketones do seem more clustered than expected by chance, but our current data are insufficient to confirm this suggestion.

**Fig. 3.**
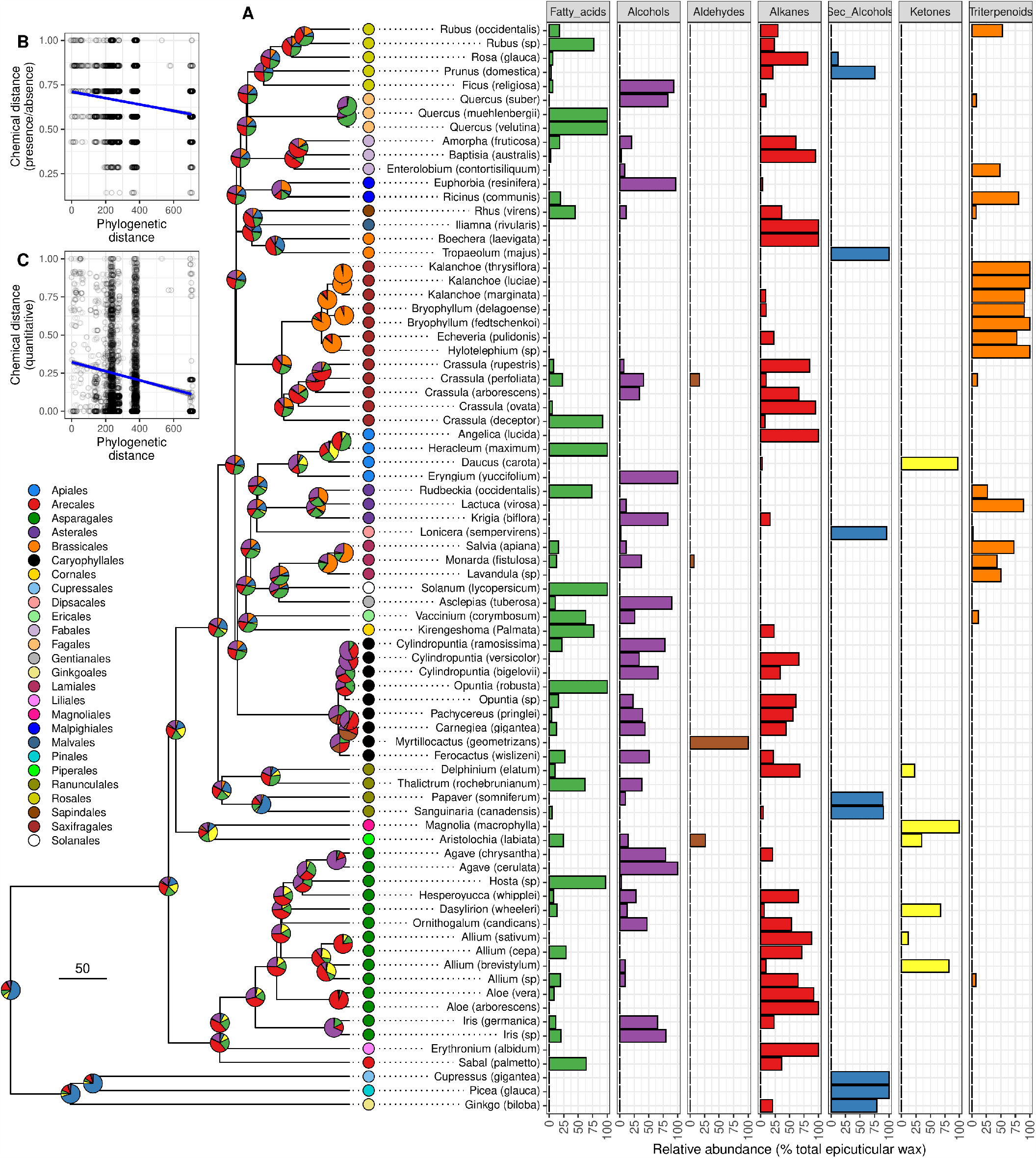
Distribution of epicuticular wax bloom compound classes across plant species. **A**. Phylogeny showing the relationships between the species sampled as part of this study. The phylogeny was derived by pruning a previously published plant megaphylogeny (27). The tips of the phylogeny are labelled with colored dots that indicate the family to which each taxon sampled belongs, as indicated in the color legend. The data presented in the accompanying horizontal bar chart were collected by (untrained) citizen scientists, which resulted in some uncertainty in the species-level identifications. Accordingly, the tips of the phylogeny are labelled with the name of the genus to which the taxa belonged, and the citizen scientits’ identifications of the species’ names in parentheses. The accompanying horizontal bar chart shows the relative abundance of each compound class detected in the epicuticular wax bloom of each species. Each compound class is given a different color. Those sample colors are also used in the pie charts on each node of the phylogeny, where each pie charts shows the estimated chemical composition of ancestral chemical blooms. **B**. Regression analysis of the proportion of compound classes shared by a given pair of species versus the phylogenetic distance separating them. **C**. Regression analysis of the quantitative similarity between the chemical profiles of a pair of species versus the phylogenetic distance separating them. In both B and C, each point represents a pair of species, and the line of best fit was drawn based on the least squares fit for all the pairs.

Our finding that secondary alcohols exhibited significant phylogenetic signal suggested that one or more relatively recent evolutionary events were responsible for the accumulation of secondary alcohols in epicuticular wax blooms. Indeed, previous work has shown that cytochrome P450 oxidases (for example, MAH1 in *Arabidopsis*) are responsible for the generation of wax secondary alcohols from alkanes (28, 29). Our data, in combination with data from previous work on P450 oxidases, suggest that the biosynthesis of secondary alcohols has evolved relatively recently compared to evolution of other wax compound classes. In contrast, triterpenoids (the other class for which we detected statistically significant phylogenetic signal) are known to be ancient compounds. They have been found in virtually every major plant lineage and in algae (30, 31). Our finding that triterpenoids exhibited statistically significant phylogenetic signal was, therefore, surprising. One possible explanation is that the biosynthesis of triterpenoid compounds is ancient, but that some other process required for triterpenoid accumulation at the plant surface is newer and only appears in some lineages. Once cuticular wax compounds have been synthesized, they are transported to the surface by a collective of transport machinery, including ABC transporters and non-specific Lipid Transport Proteins (32, 33). Thus, we hypothesize that the machinery dedicated to transporting triterpenoids to the plant surface is a relatively recent innovation that has allowed triterpenoid-rich epicuticular wax blooms to appear in statistically significant lineage-specific patterns. We applied ancestral state reconstruction to the wax chemical data presented here (Fig. 3, pie charts at tree nodes), and identified two nodes at which triterpenoids may have been highly abundant in epicuticular wax blooms. These nodes are the common ancestor of *Bryophyllum, Kalanchoe, Echeveria*, and *Halyphytum*, as well as the common ancestor of *Salvia, Monarda*, and *Lavandula* species. We suggest future comparisons of these lineages against their close relatives as a means to test the hypothesis that transporting triterpenoids to the cuticle for epicuticular wax production is a lineage-specific phenomenon.

In addition to detecting phylogenetic signal for secondary alcohols and triterpenoids on the compound class level, we detected significant phylogenetic signal for a number of individual compounds. Apart from some secondary alcohol compounds and triterpenoid compounds, we detected signal for C21, C25, and C27 alkanes, as well as C27 and C29 fatty alcohols. Alkanes in plant waxes are most typically found with odd chain lengths C27-C33 and fatty alcohols are most often found with even total carbon numbers. Thus, within their respective compound classes, these chain lengths are somewhat unusual. This suggests that even though the biosynthesis of alkanes and fatty alcohols seems ancient and widespread among extant land plants, the synthesis of specific chain lengths within those classes may be lineage-specific and thus a more recent innovation.

## 3. Materials and Methods

Citizen scientists collected wax bloom samples on cotton swabs. Once received in the laboratory, each sample was given an ID number and metadata was recorded, including the collector’s name, the state of origin, and the plants scientific and common name. Next, each cotton swab was used to prepare a GC-MS sample. To prepare a sample, a GC vial was labeled according to the ID of each sample and 1 mL of chloroform was added to each vial. Each swab was removed from its pouch and was immersed in the respective vial containing chloroform for 1 minute, then the chloroform was allowed to evaporate in a fume hood. Once dry, 125 uL of pyridine and 125 uL of N,O-Bis(trimethylsilyl)trifluoroacetamide (BSTFA) were added to the vials to convert wax compounds into their trimethylsilyl derivatives. Vials were then capped, vortexed, and finally incubated at 70 °C for 45 minutes.

Prepared GC samples were run on a 7890B Network GC (Agilent) equipped with an 7693A Autosampler (Agilent) equipped with a split/splitless injector and an HP-5 capillary column (Agilent, length 30 m x 0.250 mm x 0.25 μm film thickness). Each sample was injected with a split/splitless injector with He as the carrier gas with a flow rate of 1 mL/min. Oven conditions were as follows: 50 °C hold for 2 minutes, 40 °C/minute ramp to 200 °C, hold for 2 minutes, 3 °C/minute to 320 °C, hold for 2 minutes. MS conditions were: 70 eV EI ionization, one scan cycle per second. Since the concentration of analytes in the sample was not known, an initial injection volume of 1 uL was utilized. After a sample was run, the resulting total ion chromatogram was inspected and injections of a higher volume or concentration were conducted as necessary for each sample.

All phylogenetic analyses reported here were implemented in R. To generate a phylogeny for the 78 species for which we obtained chemical data, we pruned a previously published megaphylogney (27) so that only tips corresponding to the 78 remained. In some cases, the published phylogeny did not contain a tip for a species for which we had chemical data, so we placed that species’ name on the tip for another species in the same genus. To compute chemical distances between pair of species, two metrics were used: the proportion of compound classes shared by the two species and the Euclidean distance between the profiles of the two species. Phylogenetic distance was obtained using the cophenetic.phylo function from the R package ape (34). To conduct ancestral state estimations, the fastAnc function from the phytools packages was used (35). For phylogenetic signal, Blomberg’s K (36) was calculated using the phylosignal function from the R package picante (37).

## 4. Acknowledgements

The authors wish to acknowledge Evan W. LaBrant for assistance with initiating this project. We also acknowledge and are very grateful to all the citizen scientists who participated in collecting plant epicuticular wax bloom samples. Middle School students from McDonald County R-1 School District were recurring participants in our chemical bloom sample collection efforts and we give them special thanks. These students collected samples as part of either Mr. Keith Jones’ 7th Grade Science class, or through the school district’s SOPE program: a place-based, argument driven inquiry program Mr. Jones leads to engage students in local conservation science and ecology. We are also grateful for support from the University of Minnesota’s UROP program and the University of Minnesota Duluth’s BURST and SURP programs. Dien Nguyen acknowledges support from a Summer QMA Graduate Fellowship and the Cothran Memorial Fellowship. Lucas Busta acknowledges support in the form of start up funds from the University of Minnesota Duluth and a postdoctoral research fellowship from the National Science Foundation’s Plant Genome Research Program (NSF PRFB IOS-1812037).

## 6. Supplementary Material

- Supplemental Figure 1: **Distribution of epicuticular wax bloom compound classes across plant species**.
- Supplemental Table 1: **Chemical and Phylogenetic Data**.

**Fig. S1.**
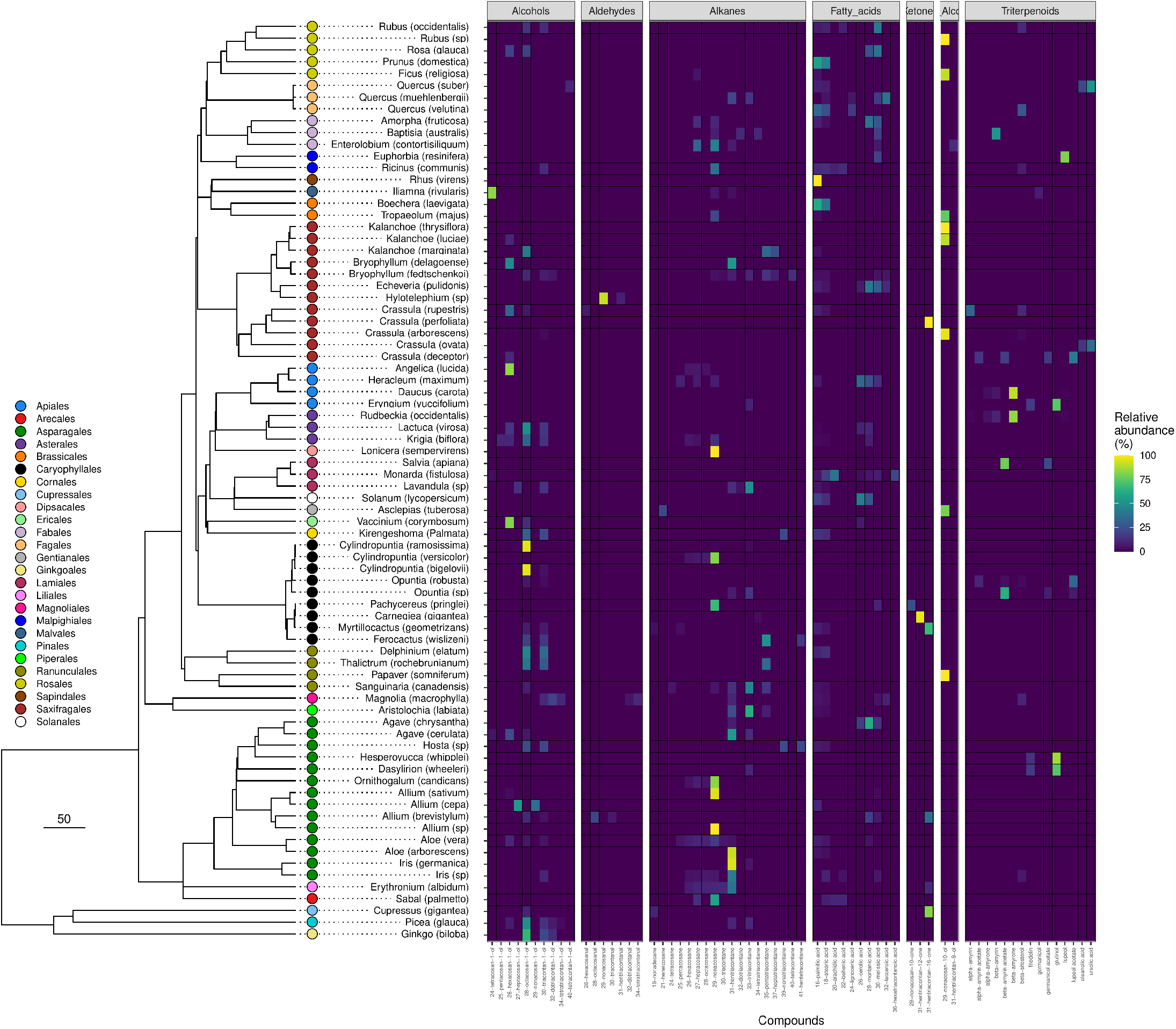
Distribution of epicuticular wax bloom compound classes across plant species. Phylogeny showing the relationships between the species sampled as part of this study. The phylogeny was derived by pruning a previously published plant megaphylogeny (27). The tips of the phylogeny are labelled with colored dots that indicate the family to which each taxon sampled belongs, as indicated in the color legend. The data presented in the accompanying horizontal bar chart were collected by (untrained) citizen scientists, which resulted in some uncertainty in the species-level identifications. Accordingly, the tips of the phylogeny are labelled with the name of the genus to which the taxa belonged, and the citizen scientits’ identifications of the species’ names in parentheses. The accompanying heat map shows the relative abundance of each compound detected in the epicuticular wax bloom of each species.

